# Encoding of locomotion kinematics in the mouse cerebellum

**DOI:** 10.1101/078873

**Authors:** Tomaso Muzzu, Susanna Mitolo, Giuseppe Pietro Gava, Simon R Schultz

**Affiliations:** Centre for Neurotechnology and Department of Bioengineering Imperial College London Exhibition Road London SW7 2AZ United Kingdom

## Abstract

The cerebellum has a well-established role in locomotion control, but how the cerebellar network regulates locomotion behaviour is still not well understood. We therefore characterized the activity of cerebellar neurons in awake mice engaged in a locomotion task, using high-density silicon electrode arrays. We characterized the activity of over 300 neurons in response to locomotion, finding tuning to speed of locomotion, turning, and phase of the step cycle. We found that the cerebellar neurons we recorded mainly encoded information about future locomotor activity. We were able to decode the speed of locomotion with a simple linear algorithm, needing relatively few well-chosen cells to provide an accurate estimate of locomotion speed. Our observation that cerebellar neuronal activity predicts locomotion in the near future, and encodes the required kinematic variables, points to this activity underlying the efference copy signal for vertebrate locomotion.

## Introduction

An animal’s survival relies heavily upon its ability to locomote through space. The ethological importance of locomotion is reflected in the large proportion of the central nervous system involved in locomotor activity. One such area is the cerebellum, whose function was long ago established through clinical and lesion studies to be essential for learning and controlling movements (Flourens, 1824; Luciani, 1891; Holmes, 1939). Being located, in circuit terms, between higher cortical centres and the periphery, the cerebellum acts as a strategic relay point for sensorimotor integration.

Electrophysiological studies combined with the analysis of behaviour provided direct evidence for the role of the cerebellum in locomotor control and learning. The spinocerebellum, the central part of the cerebellum, receives projections from the spinal cerebellar tract neurons which convey peripheral sensory signals and information from the spinal pattern generator (Arshavsky et al., 1983; Fedirchuk et al., 2013). In particular, the medial zone of the spinocerebellum, the vermis, has been identified as the area involved in controlling posture, tone, flexion and extension of limbs (Chambers and Sprague, 1955).

The spinocerebellar tracts, which are part of the locomotion circuitry (Goulding, 2009), were found to be essentially preserved across animal species, including mice (Oscarsson, 1965; Berretta et al., 1991; Sengul et al., 2015). The mouse is a model organism of particular interest due to its suitability for the use of transgenic technology to dissect out the contributions of individual circuit elements. In recent years, the application of transgenic techniques to mouse experiments provided new insights into the neural circuits involved in locomotion (Akay et al., 2014; Bellardita and Kiehn, 2015; Kiehn, 2016), and the role of the cerebellum in motor and cognitive functions (Reeber et al., 2013; Zhou et al., 2014; Galliano and De Zeeuw, 2014; Hoogland et al., 2015).

Observation of neural activity in the cerebellum has revealed that many cerebellar neuron types carry locomotion-related information. Purkinje cells, the only output of the cerebellum, are essential to interlimb coordination, adaptation to external perturbation, and they tend to fire rhythmically with the stepping cycle (Yanagihara and Kondo, 1996; Ichise et al., 2000; Orlovsky, 1972; Armstrong and Edgley, 1984). Although Purkinje cells in the cat were not observed to have substantial firing rate modulation by walking speed on a treadmill (Armstrong and Edgley, 1988), it was recently observed that the firing rate of Purkinje cells, averaged within single steps, can be modulated both negatively and positively with locomotion speed in freely running rats (Sauerbrei et al., 2015). Golgi cells were also shown to discharge rhythmically during locomotion, however no modulation by the speed of locomotion was observed in this case (Edgley and Lidierth, 1987). In contrast, granule cells and molecular layer interneurons of mice on a a spherical treadmill increased their firing rate during locomotion relative to stationary periods (Ozden et al., 2012; Powell et al., 2015), leaving open the question of whether and how cerebellar neurons are tuned to the speed of locomotion.

Electrophysiological recordings of single units in the cerebellum validated the relationship between behaviour and neural activity, but have thus far failed to account for population dynamics, providing a confined view of the cerebellar neural code. In fact, cerebellar circuitry is characterised by a high degree of feedback, feed-forward and collateral connections (Ito, 2006; Coddington et al., 2013; Mathews et al., 2012; Rieubland et al., 2014; Astorga et al., 2015), and a distinctive divergence-convergence configuration of inputs and outputs (Napper and Harvey, 1988; Person and Raman, 2012). There are therefore important unanswered questions as to how locomotion-related information is conveyed by ensembles of cerebellar neurons, and what type of neural population code is employed.

To address these questions, we recorded from movement-sensitive populations of neurons in lobules V and VI of the cerebellar vermis of mice navigating in a virtual reality (VR) environment. We characterised neurons whose activity is modulated by kinematics parameters such as locomotion speed and yaw rotation. The combined activity of these neurons linearly decodes locomotor speed with an accuracy proportional to the population size, providing new insight on the neural code of the cerebellum.

## Results

Mice (n=14, 16-24 weeks old) were head-fixed on an air-supported spherical treadmill inside a demispherical screen (Figure 1A; Holscher et al., 2005; Harvey et al., 2009). Sphere pitch and yaw movements generated by the mouse were read by two optical computer mice, and integrated in order to determine the translation and heading of the animal in the virtual space. Visual stimuli were controlled in closed-loop using custom-developed LabView software (see Methods). The mice navigated through a virtual corridor along which they received water at defined reward points (green cylinders, Figure 1B). Following behavioural training (Figure 1C and Figure 1 - figure supplement 1), multi-electrode array extracellular recordings were made from lobules V and VI of the cerebellar vermis (Figure 1D). Four animals did not receive behavioural training, but instead were allowed to run in the dark, with the virtual reality stimulus switched off, as a control group to discern the influence of visual feedback on locomotor speed encoding. Animals spent on average 57*±*3% (mean*±*s.e.m., n=39) of each recording period running (defined as speed exceeding 1 cm/s). For each animal, the recording electrode was positioned at a number of different depths (39 recordings in total, 311 units; see Figure 1E). Action potentials were detected and clustered to perform spike sorting (Rossant et al., 2016). For each identified unit, cell type was determined according to recently published classification criteria (Figure 1F and Figure 1 -figure supplement 2, Van Dijck et al., 2013; Hensbroek et al., 2014).

**Figure 1:**
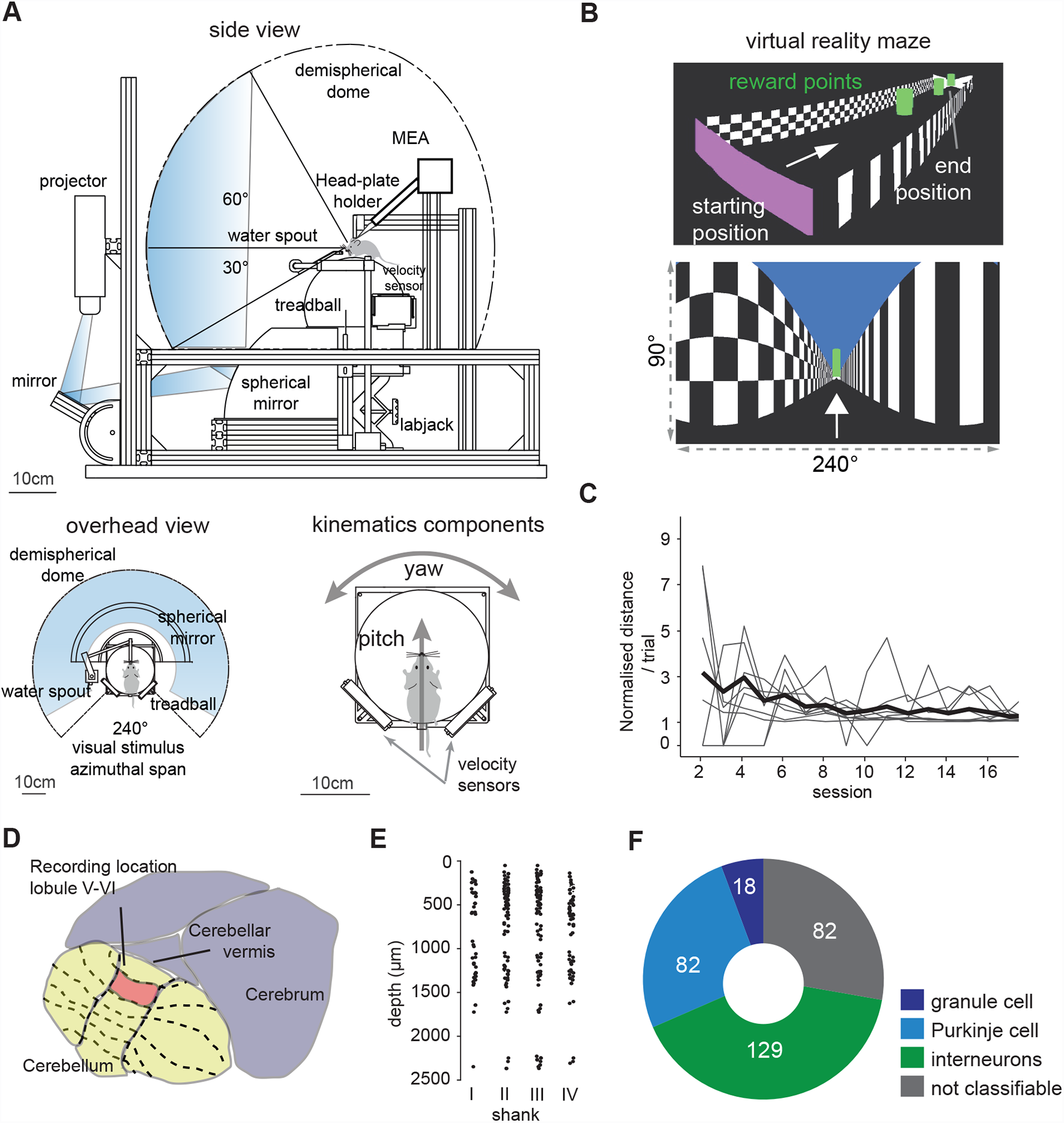
Virtual reality setup and electrophysiology recordings. (**A**) schematic view of the virtual reality system with approximate image path of the visual stimulus from the projector to the demi-spherical screen. Bottom left. Overhead view of the coverage of the mouse visual field. Bottom right. Mouse on the spherical treadmill, or treadball. The signals from the velocity sensors are integrated to determine translation (pitch) and heading (yaw) movements of the mouse in the virtual environment. Mouse drawings not to scale. (**B**) Virtual reality environment. Top, perspective view of the virtual corridor. Bottom, subject perspective of the VR maze with horizontal and vertical span of the virtual camera. (**C**) Mean normalised distance ran per trial in first 16 days. Gray lines are subjects, thick black line is average for all mice. (**D**) Recording location area. (**E**) Depth of single units (n=311) from the cerebellar surface grouped by shank (I-IV) for all mice. Depth measurements are based on the position of the channel in which the greatest spike amplitude of the signal is recorded (**F**) Pie chart of cell classes as identified by a hybrid classification algorithm based on VanDijck et al., 2013 and Hensbroek et al. 2014

### Cerebellar neurons respond to speed of locomotion

The activity of many units correlated with the behavioural status of the mouse, i.e. the firing rate changed with speed (Figure 2A-C). To determine whether neural activity correlated with speed of locomotion, we computed speed tuning curves for the firing rate of each unit (Figure 2D-F). To assess the significance of speed modulation, we shuffled the data 100 times and compared the variance of the original curve with that computed from the shuffled data. If this variance was greater than the values of at least 99 shuffled curves, the unit would be considered to be significantly speed modulated (Saleem et al., 2013; Kropff et al., 2015). We also checked whether the changes in firing rate were due only to changes in excitability between stationary and moving periods (i.e. if cells were driven by locomotion, but their activity was not modulated by speed) by repeating the above procedure but only considering speeds *>*1 cm/s. 159 units were found to respond to locomotion movements: 20 showed a binary response to movement and the remaining 139 were modulated by speed. For these units, three classes of modulation profile (tuning class) were observed: units whose firing rate monotonically increased (n=50) or decreased (n=51) with speed, and units whose firing rate reached its maximum at a preferred speed that is *≤* 70% the maximal speed achieved by the mouse (n=38). Similar profile responses were observed in na¨ıve (untrained) mice and no differences were found between the responses of units recorded from these and trained animals. These units were therefore analysed conjointly (Figure 2 - supplement figure1A).

**Figure 2:**
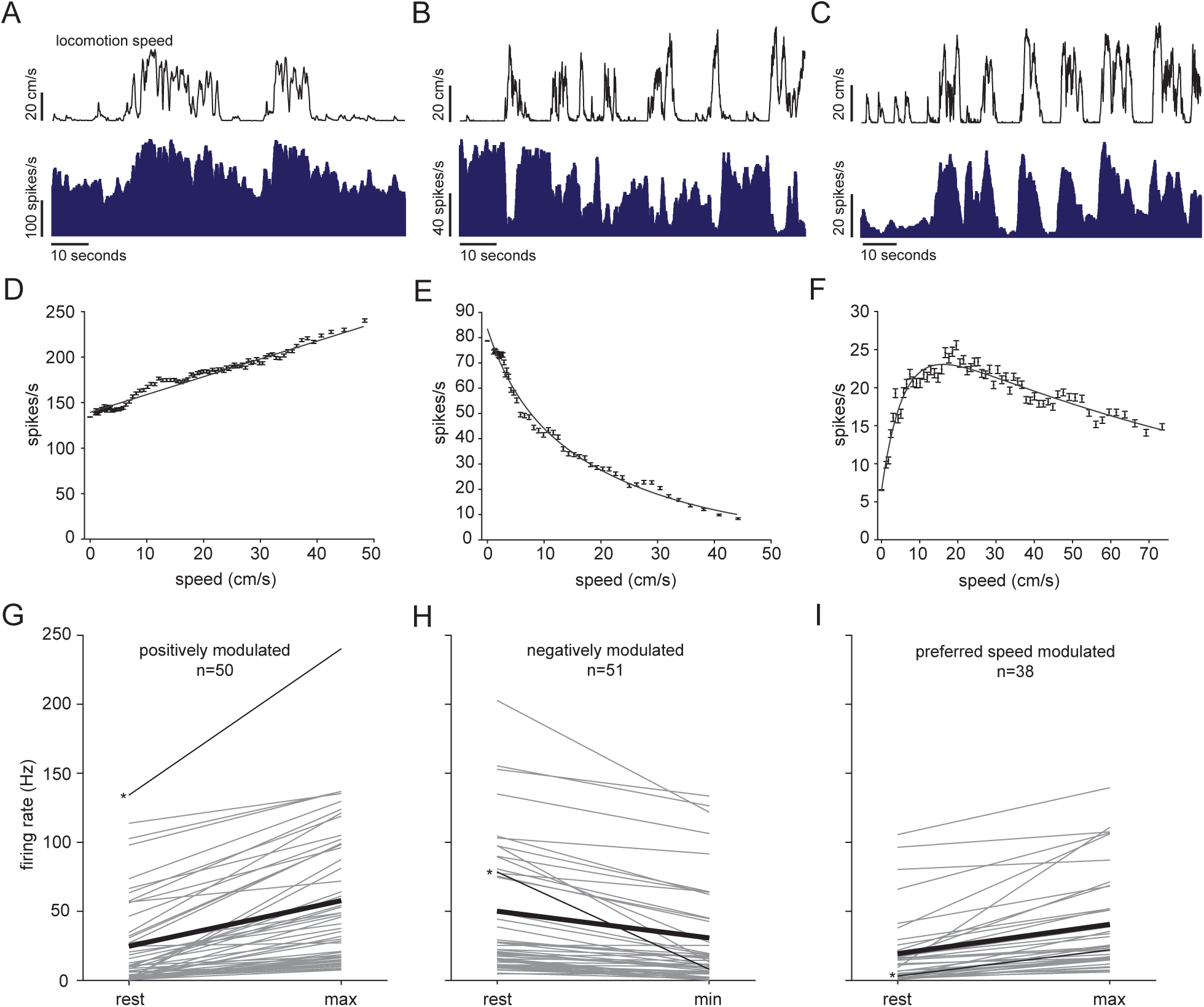
Cerebellar neurons response to locomotion speed. (**A-C**) Speed and instantaneous firing rate of three example units for the three response profiles observed (bins = 0.5 seconds). (**D-F**) Speed tuning curves of the examples in A-C; error bars are standard errors. (**G-I**) Mean firing rate during stationary periods and speeds at which maximal firing rate is recorded. Gray lines are single units; thin black lines with asterisks on the left indicate units shown in A-C and D-F; thick black lines are average firing rates of all units within each response class.

Positively modulated units, on average, had lower spontaneous (animal at rest) firing rates compared to those cells whose firing rate decreased with speed (Figure 2G,H). Firing rate changed from 24.7*±*4.6 Hz during resting periods to 52.2*±*7 Hz (mean*±*s.e.m., n=50) at maximal locomotion speed. For negatively modulated units, firing rate decreased from 50*±*7.6 Hz under the resting condition to 30*±*6.1 Hz (mean*±*s.e.m., n=51) during locomotion at maximal speed. Units showing a preferred speed had on average less marked changes going from 19.3*±*4.2 Hz at rest up to 40*±*5.7 Hz (mean*±*s.e.m., n=38) at the maximum speed observed (Figure 2I).

We examined whether the modulation of single cells differed within and across response tuning classes (Figure 2 - supplement figure 2A). The modulation index was defined as the ratio between the difference and the sum of the maximum and minimum firing rates measured on each tuning curve. Modulation indexes for all response types varied heterogeneously across the whole range. We found that, for all response tuning classes, the modulation of firing rate was negatively correlated with spontaneous firing rate (Figure 2 - supplement figure 1B).

Units responsive to movement were observed in all animals, with no discernible dependence on depth of recording site (Figure 2 - supplement figure 3). Units belonging to the same response tuning class were not observed to cluster spatially: in only 8 out of 39 recordings did we find units belonging to the same response class in close proximity (i.e. in the same electrode shank). In all remaining recordings (n=27) in which we found multiple units responsive to movement in close proximity, their response type was heterogeneous.

Taken together, these results demonstrate that cerebellar neurons respond to locomotion speed by either increasing or decreasing their firing rate or responding maximally to a particular speed. Furthermore, we did not find spatial clustering of units with the same tuning profile class. This observation suggests that nearby cerebellar neurons, possibly belonging to the same micro-zone (Oscarsson, 1979), encode locomotion-related movements by combining different types of sensorimotor information.

### A subset of cerebellar neurons display yaw direction tuned responses

While animals ran in the virtual corridor, they corrected their trajectories repeatedly to reach the target location marking the end of each trial. As the animals exerted more strength on either one or the other side of the body when turning the sphere, we examined whether this asymmetric use of limb muscles was reflected in cerebellar neuronal activity. By extracting the yaw movement information from the motion sensors, we examined how the firing rate changed with respect to sphere rotations around the clockwise (CW, negative yaw) and counter-clockwise (CCW, positive yaw) direction (example trace, top Figure 3A).

**Figure 3:**
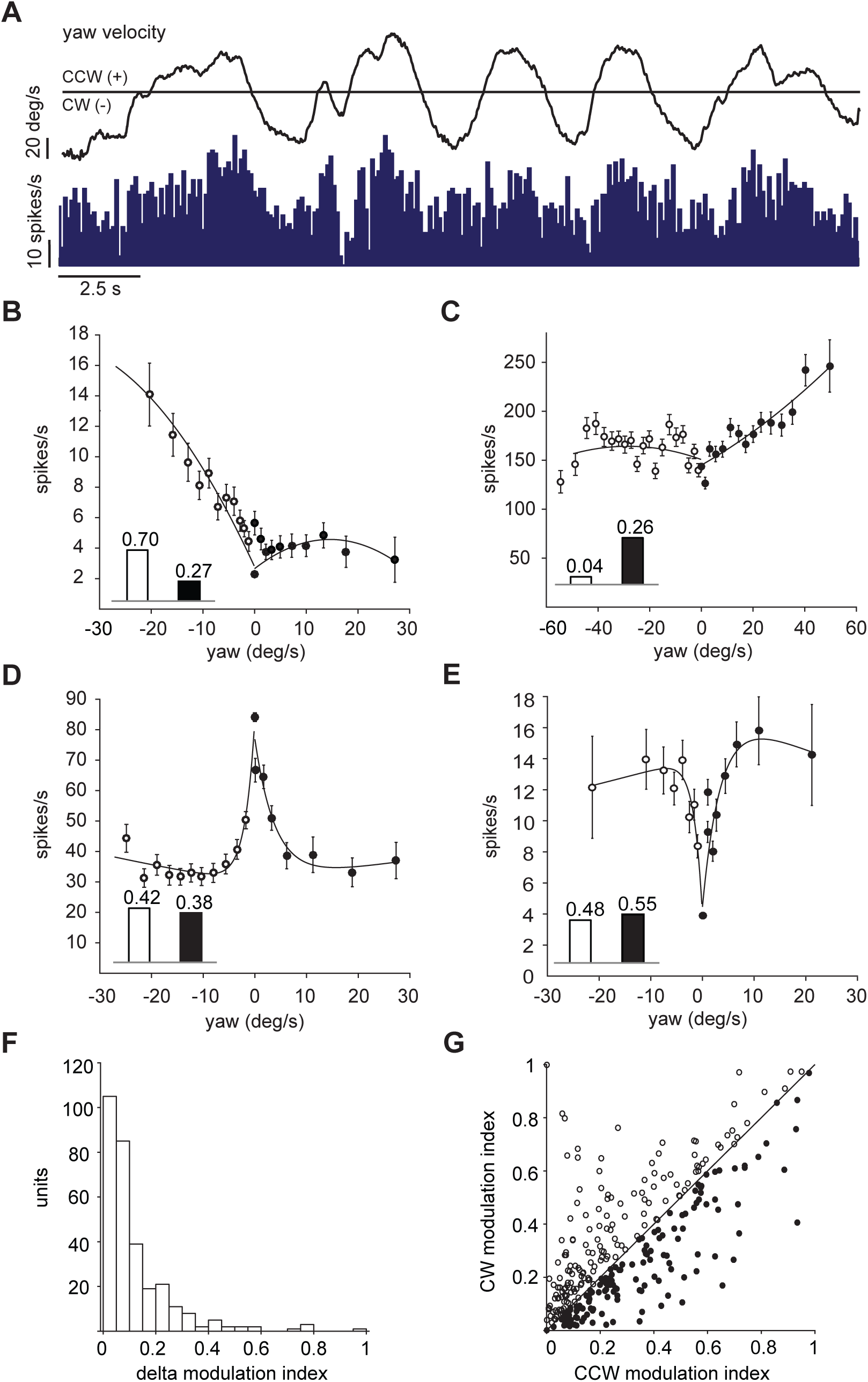
Cerebellar neurons response to yaw direction. (**A**) Example unit tuned to CCW yaw direction (positive): top, yaw trace in deg/s; bottom, mean instantaneous firing rate, bin width 100 ms. (**B**-**C**) Two example units preferentially responding to yaw-turning in the CW (**B**) or CCW (**C**); each data point is formed by more than 900 bins (100ms width); error bars are s.e.m.; inset bars indicate the modulation index values derived from the two curves. (**D**) Example unit decreasing its activity in both yaw directions. (**E**) Example unit that increases its activity in either yaw direction. (F) Distribution of the difference in modulation index between the CW and CCW yaw direction. (**G**) Population plot of modulation index values of each unit for the CW and CCW direction (n=311).

To do so, we computed tuning curves for neural activity in response to CW and CCW yaw directions, and calculated the modulation index for each cell in the two directions. For a few units, the neural response was clearly one-sided (Figure 3A). This was reflected in their tuning curves, and in the absolute difference between the modulation indexes of the tuning curves computed for the negative (CW) and positive (CCW) yaw directions (delta yaw modulation index - Figure 3B,C). Most cells, however, responded equally to either yaw direction, showing a decrease (Figure 3D) or increase (Figure 3E) in firing rate to either CW or CCW yaw movement, in an almost symmetric fashion. This is reflected in the distribution of the delta yaw modulation indexes: 63% (195 out of 311) had a difference in modulation indexes smaller than 0.1, while only 19% (58 out of 311) was greater than 0.2 (Figure 3F), indicating sensitivity to yaw direction. In fact many units had similar modulation indexes for both the CW and CCW yaw directions (Figure 3G). We also examined whether tuning to speed influenced the yaw-direction selectivity of the 58 cells that had delta yaw modulation index greater than 0.2. Interestingly, 32 out of the 58 yaw direction sensitive units did not modulate their activity with speed (Figure 3 - supplement figure 3A). There was also no evident relationship between the speed modulation index and delta yaw modulation index (Figure 3 - supplement figure 3B). These results suggest that there are units in the cerebellum that respond selectively to the use of one side of the body in preference to the other, and that these cells are not necessarily influenced by changes in speed.

### A subset of cerebellar neurons are tuned to phase of stepping cycle

Previous studies of the role of the cerebellum in locomotion control found that Purkinje cells are rhythmically modulated by stepping cycle (Orlovsky, 1972; Armstrong and Edgley, 1984; Armstrong and Edgley, 1988; Edgley and Lidierth, 1987; Powell et al., 2015; Sauerbrei et al., 2015). We reasoned that one way to produce speed tuning would be for units to respond by spiking at a fixed phase per step cycle, thus producing higher firing rates the more step cycles occur per unit of time. We therefore examined whether this was the reason behind the tuning of activity to locomotion speed. Since pitch velocity is measured by motion sensors parallel to the vertical axis, with velocity sampled at a high polling rate (f=200 Hz), it was possible to detect the vertical oscillations caused by the mouse stepping on the sphere (Figure 4A). The high frequency periodicity of the signal was extracted from the original pitch velocity signal by transforming the pitch velocity with the Hilbert operator. At least 393 putative step cycles were found for each recording session (1724*±*158, mean*±*s.e.m., n=39). We then measured the correlation between the firing rate (bin width = 5ms, smoothed with a 20-ms Gaussian window) and the phase of the step cycle for each unit (Figure 4B) finding that only 57 units out of 311 showed a significant modulation with the stepping cycle (p*≤* 0.001, *χ*^2^ test for uniformity). Of these, thirtytwo units were also significantly modulated by speed. Mean preferred phases of the modulated units were distributed across the stepping cycle, with approximately two-thirds of the response covering on average 2.2*±*0.06 (mean*±*s.e.m, n=57) radians, as measured by the standard circular deviation (Drew and Doucet, 1991) (Figure 4B).

**Figure 4:**
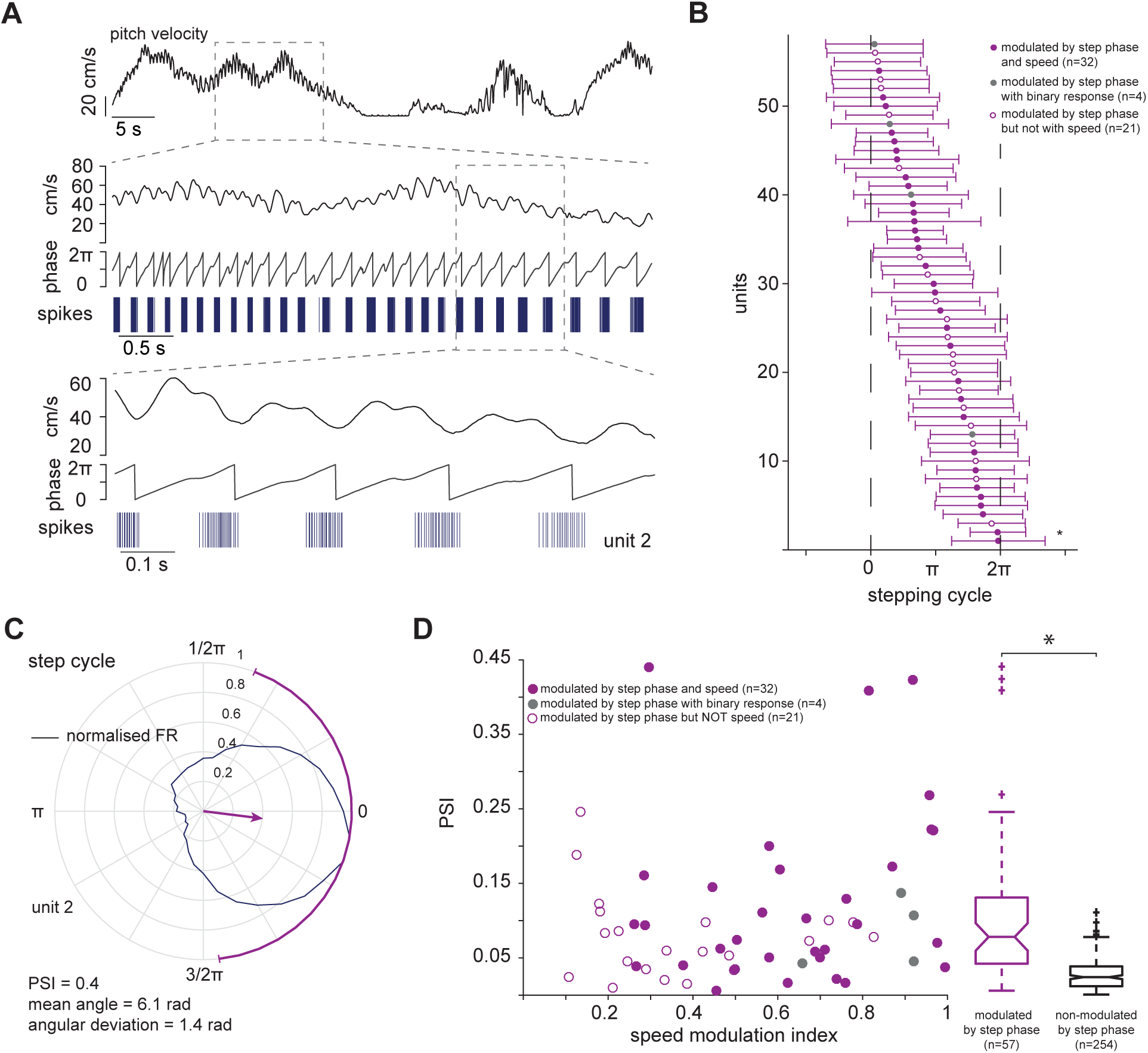
Cerebellar neurons response to stepping cycle. (**A**) Top: pitch velocity signal; centre: boxed pitch velocity signal aligned with its Hilbert transformed signal (middle) and spike events of an example unit; bottom: close-up of approximately one second. (**B**) Preferred phases of units significantly modulated by stepping cycle (n=57); horizontal bars indicate standard circular deviations. Unit 2, shown in A and C is marked by the asterisk. (**C**) Polar plot showing the preferred locomotion phase of normalised firing for the same unit shown in A; arrow indicates mean angle direction and phase selectivity index, PSI (magnitude of mean response); purple solid line around the circle is the circular standard deviation. (**D**) PSI of units modulated by step phase as a function of speed modulation index. Right, distribution of PSI of these units is significantly different from the distribution of the other units.

To quantify the step phase modulation of the response, we used an approach commonly used in the study of orientation tuning, an analogous problem (where here phase within a step cycle replaces orientation within a circular stimulus space). We calculated the normalised phase orientation vector and computed the orientation selectivity index (Mazurek et al., 2014), renamed here the phase selectivity index, PSI. Units significantly modulated by step phase have higher phase selectivity indexes in comparison to non-modulated units (p=2e^*−*16^, Mann-Whitney U-test, Figure 4D). These results suggest that cerebellar neurons’ activity encodes kinematic information, i.e. locomotion speed is not a by-product of rhythmic modulation of the stepping cycle as shown in a previous study (Sauerbrei et al., 2015).

### Cerebellar units compute multiple kinematic parameters

We have described units tuned for speed of locomotion, yaw (including some tuned for direction of yaw motion), and phase of stepping cycle. It is important to determine whether these constitute separate classes of neurons, or if instead, each of the neurons displays tuning to a lesser or greater extent across each dimension.

For each cell, we therefore compared the modulation index for speed with the mean modulation index for CW and CCW yaw (see Methods), and with the phase selectivity index. These are depicted in a tri-plot in Figure 5A. It is apparent that speed, yaw and phase selective units do not cluster in this space, but are instead distributed relatively uniformly. A similar picture arises when instead comparing the mutual information units convey about speed, yaw and step phase (Figure 5B). As speed and yaw are not independent - indeed, speed is comprised of both a pitch and a yaw component - we broke speed tuning up into these components and assessed them separately. Figure 5C shows that units that conveyed substantial information about pitch also tended to convey substantial information about yaw - and that in fact, more of the speed information arose from yaw than from pitch signals. As bimodal distributions were not found in the information conveyed about speed, pitch, yaw nor step phase, we conclude that the cerebellar units examined were a relatively homogeneous population encoding all of these quantities to a greater or lesser extent, rather than a heterogeneous population comprising clusters of units encoding different kinematic variables.

**Figure 5:**
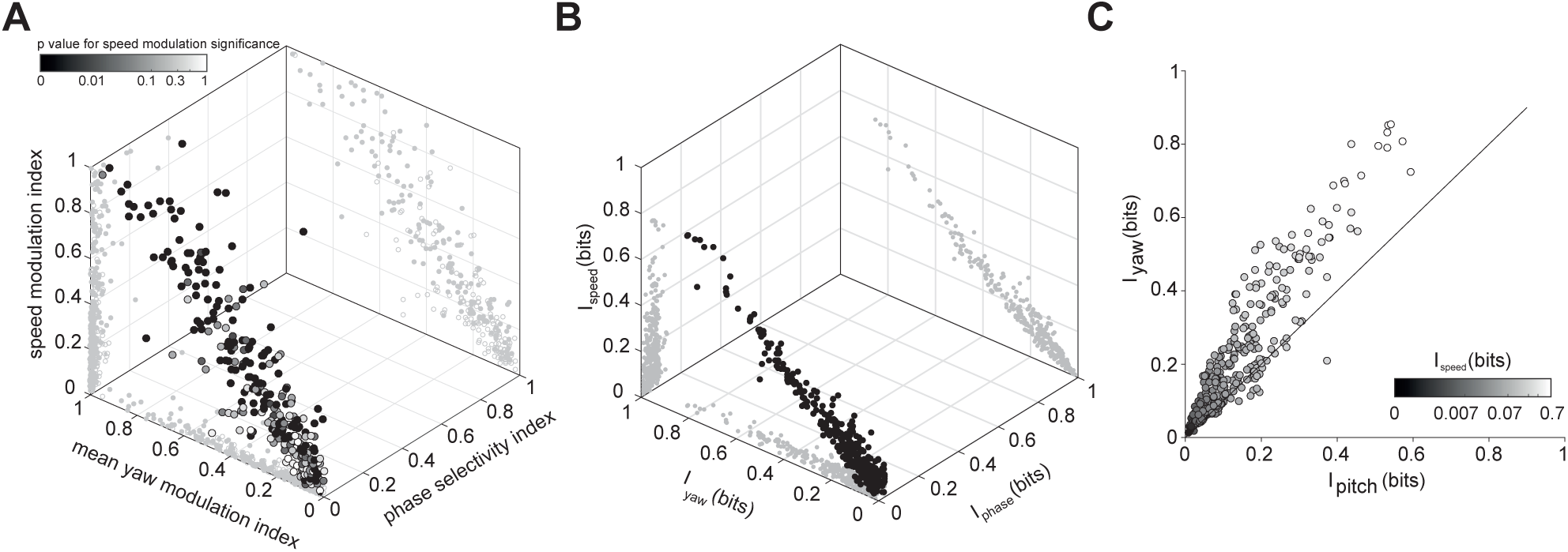
Modulation indexes and mutual information values of cerebellar units. (**A**) Modulation indexes for speed, yaw and phase selectivity index. Colour coding indicates p value of test for significance of speed modulation. Mean yaw modulation index is the mean between the modulation indexes of the yaw tuning curves for clockwise and counter-clockwise directions. (**B**) Mutual information values for speed, yaw and step cycle. (**C**) Mutual information values of the two vectorial components of speed, pitch and yaw. Colour coding indicates the value of mutual information of firing rate and speed.

### Cerebellar neurons mainly encode motor information

In order to understand whether the recorded units encode information about descending motor commands, we analysed the timing of the information conveyed by neuronal responses about locomotion speed, reasoning that for motor units, the firing rate should provide predictive information about locomotion speed, whereas for sensory units, the information should be largely retrospective. We computed the mutual information (Schultz, Ince, et al., 2015) between the firing rate and locomotion speed time courses for each neuron, for a range of imposed time lags. Firing rates were shifted with respect to the speed signal by 10 ms from -500 to +500 ms (Figure 6A). While there were retrospective units (peak mutual information at negative time lag), the majority of speed modulated units showed a peak in the mutual information at a positive time lag of 108.1*±*17.2 ms (mean*±*s.e.m., n=139), suggesting that the neurons observed may primarily encode descending motor signals rather than sensory feedback (Figure 6B).

**Figure 6:**
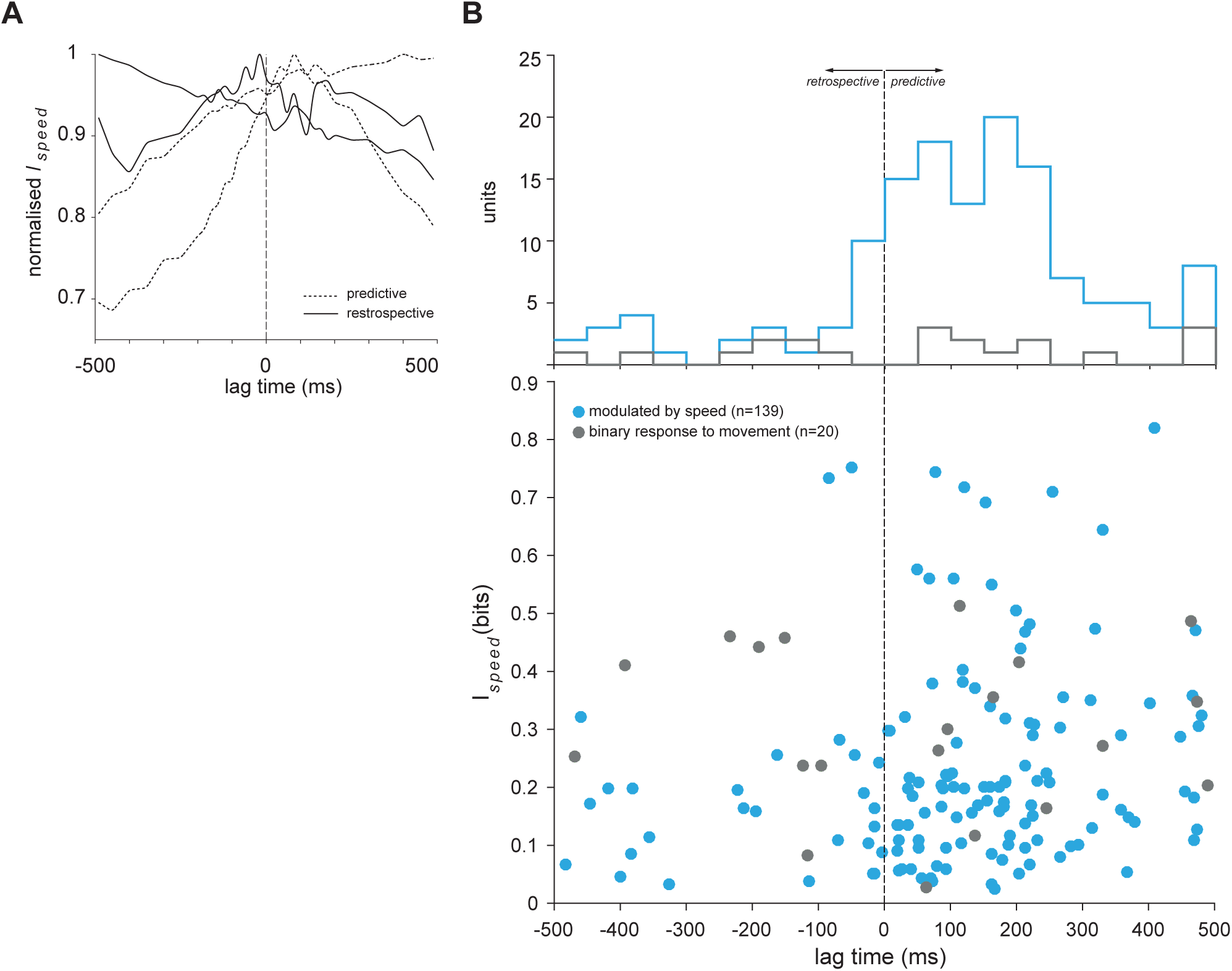
Mutual information of locomotion speed and firing rate. (**A**) Normalised mutual information of four example units for different time lags; predictive units are shown in dashed lines and retrospective units in solid lines; for each type, two examples are shown: one that either increases or decreases with time and one with a peak either before or after zero lag. (**B**) Top: Histogram of the distribution of the peaks at which maximal mutual information is found for units responsive to movement and modulated by speed (bin size = 50 ms). Bottom: maximal mutual information values of the units responsive to movement and modulated by speed with respect to the time lag.

The values of maximal mutual information were significantly higher that those of non-modulated units (p=2.3e^*−*9^, Mann-Whitney U-test): 0.24*±*0.01 bits (mean*±*s.e.m., n=139) for speed modulated units and 0.13*±*0.01 bits (mean*±* s.e.m., n=172) for unresponsive units. In fact, the lower information values - calculated at zero lag - for the latter class are consistent with the classification based on the tuning curve modulation approach (Figure 6 - figure supplement 1).

### Speed of locomotion can be linearly decoded from cerebellar neuronal ensembles

It has been previously shown that neural activity of single Purkinje cells encode multiple kinematic parameters of multi-joint movements during arm reaching tasks in primates (Roitman, 2005; Pasalar et al., 2006; Hewitt et al., 2011). Similarly, locomotion requires the coordination of multiple joints and muscles. Since the majority of units we found was shown to encode multiple kinematics parameters, we set out to find out whether locomotion speed could be accurately reconstructed by populations of cerebellar neurons.

To this end, we developed an optimal linear estimator (OLE) with the aim to reconstruct the locomotion speed time course from a weighted sum of the firing rates of each unit (Figure 7A). The weights were adjusted by linear regression, so as to minimise the mean squared error (MSE) between the decoded and the original recorded trace. The decoder was trained on 70% of the data of each recording and tested on the remaining 30%. To assess the scaling of decoder performance with ensemble size, we selected recordings comprising at least eight units (n=6).

**Figure 7:**
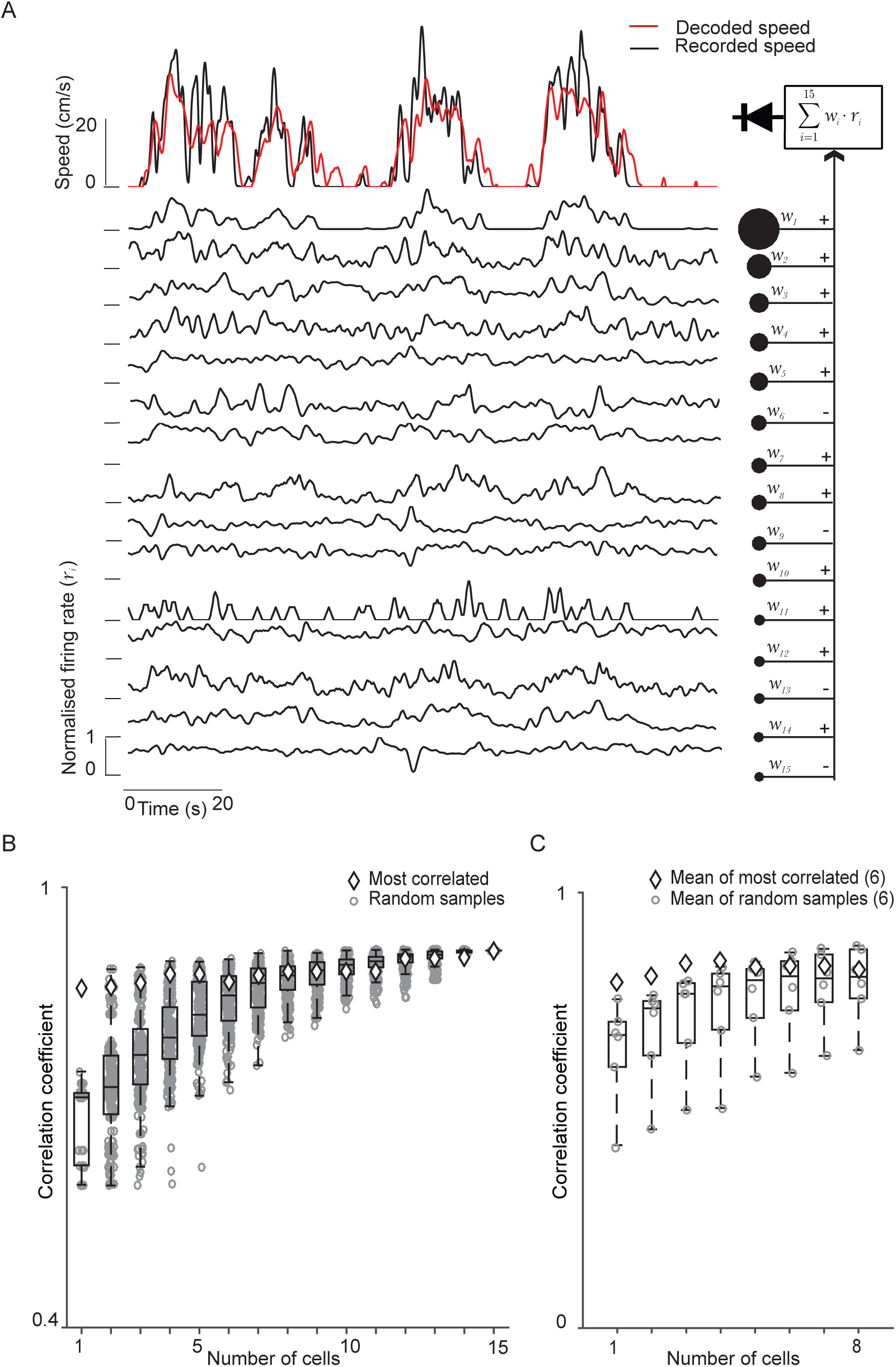
Population decoding of cerebellar neurons reconstruct speed. (**A**) Diagrammatic representation of the optimal (least MSE) linear decoder used to reconstruct locomotion speed showing one example experiment with 15 units. Top: Time course of original and reconstructed locomotor speed signal. Bottom, normalised firing rate of all units used for the decoder. Right: firing rates of all units are weighted and linearly summed with the result then being half-wave rectified (diode symbol). (**B**) Reconstruction quality as a function of the number of the units used for the decoder of the same example experiment shown in A. Box plots show first and third quartile with middle horizontal line indicating the median of 2*N* combinations of populations’ accuracy. (**C**) Mean reconstruction quality as a function of the number of the units for 6 different experiments. The average of the most correlated ensembles is plot together with the mean of the randomly formed ensembles.

We investigated how the population size affected the accuracy of the decoder. To avoid any bias in the choice of units used in the reconstruction, these were selected randomly for any given size of neural population. 2*N* possible population combinations were tested for ensembles of 1, 2, *N −* 1 and *N* units (*N* number of units recorded in the experiment), and *N* ^2^ otherwise. As the number of units increased, so did the decoder accuracy whilst the decoding performance range of the random selected ensembles reduced (Figure 7B). Furthermore, as the population size increased, the median accuracy of locomotion speed reconstruction approached the one obtained using populations formed by the most correlated units only, as measured by the Pearson correlation coefficient.

These results are consistent across all experiments that contained more than seven units (Figure 7C) suggesting that the cerebellum encodes kinematic information related to instantaneous locomotion speed by linearly summing the contributions of single neurons.

## Discussion

In this study, we used a virtual reality behavioural task, together with multi-unit electrophysiological recording, to investigate the neuronal population activity underlying locomotion. Our approach allowed us to assess the activity of multiple neighbouring neurons during behaviour while maintaining a high degree of experimental control over behavioural parameters. Our recordings from lobules V and VI of the vermis indicate that most cells in this area encode kinematic parameters of locomotion. We found that while 6% of the cells showed significantly different firing rates during locomotion as opposed to rest, but no significant modulation by speed (similar to the classical results of Armstrong and Edgley (1984)), 45% of neurons were specifically tuned for speed of locomotion. This included cells that increased their firing rate with increasing running speed, cells that decreased their firing rate below the spontaneous (rest) level with increasing speed, and cells showing a preferred locomotion speed. We also found that some cells were tuned to yaw (turning) and also to particular phases of the locomotion step cycle. The different responses of individual cells that we observed to locomotion speed, turning and stepping may reflect interdependent information about behavioural state, laterality and type of muscles being exerted during locomotion.

While we have employed and extended recently developed cell classification algorithms (Van Dijck et al., 2013; Hensbroek et al., 2014), our approach is limited by the lack of certainty about cell identity. We were able to identify Purkinje cells, granule cells, and interneurons, however we were not able to classify interneurons into specific classes with any degree of confidence, and our approach did leave a significant number of unclassified neurons. In the present study, for technical reasons we did not attempt to analyse complex spike (CS) waveforms from Purkinje cells, restricting our attention to simple spike (SS) waveforms. CS will be the subject of a future manuscript. Further improvements to cell classification algorithms will probably require obtaining ground truth validation data through simultaneous MEA and sharp electrode or whole cell patch clamp recording in the awake animal, with histological validation, a technically challenging task. In our study, however, we did not find the encoding of locomotion kinematic parameters to be dependent upon cell class.

Kinematic parameters of arm movements have been found to be related to single neuron activity in the cerebellum (Hewitt et al., 2011; Popa et al., 2012) and in the motor cortex (Ashe and Georgopoulos, 1994; Yu et al., 2007) of non-human primates. It has recently been reported that Purkinje cells discharge rhythmically during locomotion (Sauerbrei et al., 2015), and that granule cells fire in bursts at locomotion onset (Powell et al., 2015). Ozden et al. (2012) reported finding molecular layer interneuron activity to increase following transition from a resting to a moving (locomotion) state. Sauerbrei et al. (2015) did report modulation by speed of some of the cells in their dataset (recorded from the freely moving rat), but did not describe speed tuning further, instead focusing on step-phase dependent correlations with behaviour. Moreover, vestibular inputs were not taken into account in their modulation analysis; such inputs were controlled in our study, as we used head-fixed animals for which these inputs can be considered negligible. To the best of our knowledge, the current study is the first description of the tuning of cerebellar neurons to speed of locomotion. One question that may arise is why this was not observed previously, for instance in earlier studies of cats walking on a treadmill. While it is not at this stage possible to rule out that inter-species differences account for this, our view is that the discrepancy is more likely to arise from the fact that in these studies the cats passively stepped on a treadmill, which was rotating at a rate fixed by the experimenter. Instead, in our study, the animals actively locomoted at a speed of their own choice (starting and stopping as they wished), motivated by an increasing reward rate for more rapid progress down the corridor segments, which may engage cerebellar networks to a greater degree. In fact, a small amount of speed modulation is apparent in Fig. 2 of Armstrong and Edgley (1988) for most of the neurons they recorded, suggesting that the situation in the mouse and cat may not be completely dissimilar.

By allowing the spherical treadmill to rotate along any orientation (rather than constraining it to a single axis as with a treadmill), we aimed to create a navigation paradigm congruent with a real-world scenario. Animals had to intentionally engage many muscles relevant for locomotion, and were not constrained to run at fixed speeds. However, the locomotion task should be thought of as a sensorimotor control task, rather than as normal locomotion behaviour, because of the artificial nature of the head fixation and of the act of balancing on a frictionless ball, which is itself an acquired skill. Because the animals were head-fixed, we assumed that vestibular inputs were negligible during yaw (turning) movements. The correlation of neuronal activity with the direction of movement may be related to lateralised spinocerebellar inputs from muscles employed to steer clockwise or anti-clockwise. We did not find cells with a preference for a turning direction (i.e. a high Delta modulation index) to be highly modulated by speed, suggesting that speed and direction locomotion information are relayed separately. A similar result was observed in macaque monkeys performing a visually guided tracking task (Roitman, 2005): Purkinje cells were found in that study to respond to position and direction of arm movement but not to arm speed.

According to our information theoretic analysis, the majority of the units provided maximum information about the speed of locomotion a short time in the future (*∼*100 ms). They can thus be thought of as providing predictive, rather than retrospective, information about locomotion, suggesting that they may be driven by internally generated rather than sensory signals. The cerebellum receives projections from the nuclues cuneiformis (Gioia and Bianchi, 1987) that, in turn, receives inputs from the mesencephalic locomotor region (MLR; Ryczko and Dubuc, 2013). Since the MLR is a region of the hind brain that is involved in initiating and modulating locomotion (Shik et al., 1966; Lee et al., 2014; Kiehn, 2016), the cerebellum might receive a copy of the motor signals sent to spinal locomotion centres (Orlovsky et al., 1999). Indeed, MLR neurons have been observed to show similar speed tuning profiles to those reported here (Lee et al., 2014). We see this in agreement with computational theories, based on forward internal models, according to which the cerebellum uses an efference copy to compensate for slow sensory feedback during fast movements (Wolpert et al., 1995; Pasalar et al., 2006). In addition, the cerebellum might use the efference copy to suppress sensory feedback in order to reduce motor noise during movements (Shergill et al., 2003; Kennedy et al., 2014; Laurens et al., 2013).

We were able to reconstruct locomotion speed to high accuracy by linearly summing (positive and negative) weighted firing rates. The performance of our decoder increased with population size, as it can be expected, suggesting that multiple motor commands and copies of the central pattern generator signals can be effectively combined to minimise noise in the output to the deep cerebellar nuclei (Eccles, 1973; Schultz, Kitamura, et al., 2009), while preserving the remarkable pattern recognition capacity of the individual Purkinje cell (Marr, 1969; Albus, 1971; Barlow, 2002). This strategy could provide a more accurate estimate of the real speed of the animal, and, in turn, optimise motor control, similarly to what has been described for population decoding of saccades duration in monkeys (Thier et al., 2000). It was striking, however, how few cells needed to be combined in order to obtain an accurate readout of locomotion speed, for well-chosen cells. This interpretation also agrees with the idea that a linear summation of different response contributions could underpin the cerebellar neural code (Walter and Khodakhah, 2009).

In this study we recorded from neurons in cerebellar vermis lobules V and VI whose activity conveyed information about locomotion kinematics. Other areas (such as paravermal lobule V, Sauerbrei et al., 2015) have also been found to represent locomotion signals. Because of the peculiar fractured somatotopy and modular organization of the circuitry (see e.g. Apps and Hawkes, 2009), it is unclear how kinematic information processing is divided across zones and how their outputs are integrated by deep cerebellar nuclei to generate motor control. Perhaps such motor estimations are also relayed to, and exploited by, other nervous centres, to integrate behavioural information relevant for motor control and spatial navigation. In future work we hope that this may be elucidated.

## Materials and Methods

### Virtual reality system

Experiments were performed in a custom made virtual reality system for mice similar to the ones used in previous studies (Harvey et al., 2009; Schmidt-Hieber and Häusser, 2013). Mice ran on a polystyrene sphere of 20 cm in diameter free floating on a 3D-printed concave inset. The motion of the sphere was read by two USB laser mice (Razer Imperator, Razer Inc, USA) positioned ninety degrees apart on the equator of the sphere. The signals carrying the instantaneous velocities of the sphere were polled at 200Hz by the host computer (Windows 7 OS, Microsoft Corporation, USA) via a Labview custom software (National Instruments Corporation, USA). These were then integrated to update the position in the virtual environment. The virtual reality environment was rendered with the openGL API implemented in the C++ language and interfaced with Labview control software via Microsoft dynamic-link libraries (DLL). This was projected via a digital projector (PJD6553w ViewSonic Corporation, USA) onto a demispherical screen around the mouse (Talbot design Ltd, UK) with a refresh rate of 120Hz. The VR apparatus comprised an automated reward system for water delivery and air puff stimulation controlled via a data acquisition card (NI BNC-2090A and NI PCIe-6321, National Instruments Corporation). Water rewards were delivered to the mouth of the mouse via a custom made water spout connected to a peristaltic pump (Model 80204-0.5, Campden Instruments). Low pressure air jets were puffed to the mouse trunk from two lateral copper tubes upon opening of two normally closed solenoid valves (model PU220AR-01, Shako Co. Ltd) connected to an air pressure regulator.

Some experiments were run in the dark by disabling the projection of the visual stimulus and only recording mouse movements on the sphere.

### Surgical procedures

All experiments were performed in accordance with the regulations of the United Kingdom Home Office. Of the fourteen animals used, ten of these animals received behavioural training before the electrophysiological recording. For the remaining four, the head stage implant and recording preparation was performed in a single surgery (as these animals did not receive any training they are referred to as ‘na¨ıve’ in this paper). All mice were implanted with a metal plate for head fixation attached on the anterior cranial bone with histoacryl tissue adhesive (Williams Medical Supplies, product code D569).

Trained mice underwent a second surgery upon completion of the behavioural training. Water restriction was terminated at least 24 hours before the surgery to make sure the animals could undergo the surgery fully hydrated. A craniotomy and durotomy were made over the central part of the vermis. The craniotomy was covered with a 1.2% agarose in with phosphate-buffered saline (PBS) and then capped with Kwik-Cast sealant and nail varnish to preserve the brain tissue until the recording session. This happened at least 24 hours after the end of the surgery.

### Behavioural training

Following a recovery period of at least 24 hours from the surgery, the mice were head-fixed on the spherical treadmill for up to 10 minutes to habituate to the system for two consecutive days. Water restriction began on day 2 post-surgery. From day 3, the virtual reality projection was switched on. This is the first session of behavioural training. To incentivize the mice participation to the task, a water reward was given through a lick port as the mouse walked underneath cylinders suspended over the corridor. The water reward was given only at the end of each trial, once the mouse reached the following two criteria: 1) when the mouse was observed to intentionally stop only to lick its water reward and 2) when the mouse could reach the end of the virtual corridor without running more than twice the length of the corridor throughout the trials. Corridor length was progressively increased up to 500 cm depending on the mouse performance and motivation. Two lateral air-puffers were used to prevent the mouse from hitting the virtual walls of the corridor. They pointed to the rear part of the trunk of the mouse on either side.

One to two hours after the end of a training session we gave the mice water *ad libitum* until they stopped drinking. Mouse weight was monitored daily as to guarantee that the each animal did not lose more than 20% of its pre-training body weight.

Mice were trained every day for at least 2 weeks and until they were capable of running 20 consecutive trials in under 30 minutes from the start of the session for two consecutive days.

### Electrophysiological recordings

After 24 hr recovery from the second surgery for trained mice (or from the first surgery for na¨ıve mice), the mice were head-fixed on the spherical treadmill, the craniotomy was exposed after gently removing all layers of nail varnish, Kwik-Cast and agarose. The electrode was inserted at a 45 degree angle along the coronal plane and allowed to stabilize in the cerebellum for approximately 10 minutes once good spike signals were detected. To record activity from lobules V and VI of the cerebellum, we used 4-shank, 32-channel multielectrode arrays (MEAs) (Neuronexus Technologies, USA, probe model Buzsaki32). Behaviour and electrophysiological activity were then recorded in parallel while the mice navigated in the virtual reality environment.

To maximise mechanical and electrical stability during the electrophysiological recording, the water spout was removed from the mouth of the mouse and airpuffers were diverted from the mouse trunk. We did not observe behavioural changes, as the mice were fully hydrated beforehand, and the airpuff noise was a stimulus strong enough to elicit trajectory corrections. Multiple recordings were acquired from each mouse (minimum duration 270 seconds, max duration 1833 seconds). The entire recording sessions lasted up to 50 minutes.

### Data analysis

#### Spike sorting and clustering

Electophysiological data from each shank were processed independently with the *SpikeDetekt* / *KlustaKwik* / *Klustaviewa* software suite (Rossant et al., 2016). After the spike times were detected and sorted with the automated program, we ‘cured’ the outcome of the spike sorting with the built-in *KlustaViewa* program. At this stage, we selected the units with the highest clustering quality. We kept units that had a central portion of the auto-correlogram completely clean (Harris et al., 2000). We merged units that were separated due to shift of the signal in different channels with the help of the clustering features viewing tool. These were also validated against each other by means of the cross-correlograms. No distinction between simple and complex spikes were considered as the time window within which each spike waveform was extracted was only 2 ms.

#### Cell classification

We used a hybrid cell classifier based on recently published algorithms (Van Dijck et al., 2013; Hensbroek et al., 2014). The algorithm was applied only to those units from which it was possible to collect at least 60 inter-spike intervals (ISI) taken during periods of stillness (speed of mouse *≤* 1 cm/s). The first steps aimed to identify putative Purkinje and granule cells based on their mean spike frequency, entropy of ISI distribution and logarithmic coefficient of variation of ISIs (CVlog), (Figure 1 - figure supplement 3); the remaining units were considered to be putative interneurons.

#### Tuning curves

Firing rate was calculated at the same frequency as the speed (5 ms bins) and then smoothed with a 150 ms Gaussian filter. Data points for speeds greater than 1 cm/s of the tuning curves are comprised by 2000 bins. The data points for speed=0 cm/s are formed by all bins taken when the mouse is still. To evaluate the significance of a unit’s firing rate modulation with speed of locomotion, we randomly shifted the spike times one hundred times by at least 20 seconds and up to the duration of the recording minus 20 seconds (Kropff et al., 2015). For each iteration, firing rate was calculated and a speed tuning curve computed, and its variance was measured. We then compared the variance of the original speed tuning curve with the ones from the shuffled data. If its value was greater than 99% of the shuffled data values, then we considered the unit as significantly sensitive to movement (binary response). We repeated this calculation and applied the same criteria to speeds *≥* 1 cm/s to assess if each unit was significantly modulated by locomotion (Saleem et al., 2013).

A unit response type was defined according to the curve that best fits the original data points. Because of the different response profiles obtained from the original data, three different curves were fitted (linear, second-degree polynomial and double exponential). The inverse of the variance of each data point was used as weight for the fitting to compensate for the different number of data points in each bin at speed=0. The coefficients of the best fit curve were used to determine the response type. In addition, we classified a cell as:

- positively modulated if the maximum firing rate was greater than the firing rate during stationary periods, and this was recorded at a speed greater than 70% of the maximum speed of the mouse;
- negatively modulated if the minimum firing rate was smaller than the firing rate during stationary periods, and it was recorded at a speed greater than 70% of maximum speed of the mouse
- having a preferred speed if the maximum firing was greater than the firing rate during stationary periods and this was recorded at a speed smaller than 70% of the maximum speed of the mouse. The tuning curves for yaw movement were calculated similarly for clockwise (CW) and counter-clockwise (CCW) turning of the sphere. We then fitted three different curves (linear, second-degree polynomial and double exponential), selected the best fitting, and calculated the modulation index for either yaw direction. Modulation indexes were calculated as:

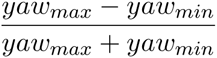

We also calculated the difference in Modulation Index (Delta Modulation Index: Modulation Index CW - Modulation Index CCW) between the CW and CCW direction to assess the ‘asymmetry’ of tuning curves. Cells with a Delta larger than 0.2 were apparently asymmetric on visual inspection.

#### Naïve vs. trained

In order to verify whether the virtual reality affected the responses to movement kinematics, we compared the population of units acquired from na¨ıve (n of units = 58) or trained (n of units = 253) animals. We used the Mann-Whitney Test to compare the modulation index distributions in the two conditions for speed and yaw. We also compared the difference in yaw modulation index and the phase index.

#### Step cycle modulation

To look at the modulation with stepping cycle, the pitch velocity signal was high-pass filtered at 3 Hz to cancel the locomotor related changes of speed. The Hilbert transform was then computed and its phase was extracted as a function of time. To ensure that pitch velocity changes were due to stepping, only putative stepping cycles longer than 50 ms and occurring only during moving periods (speed *≥* 1 cm/s) were considered. Each cycle duration was normalised with respect to time and divided in 36 equal intervals. For each interval, the instantaneous firing rate was computed.

Because of the binning of each cycle, step phase modulation was tested for uniformity with the *χ*^2^test of uniformity (Fisher, 1995). The mean direction *θ* (in radians) of the firing rate distribution of a cell around the step cycle was computed as:

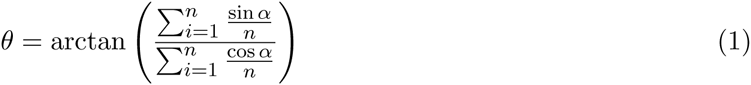

where the numerator and denominator are the mean rectangular coordinates of the resulting phase angle, *X* and *Y* respectively, *α* is the phase angle of the resultant vector 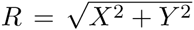 for each cycle, and *n* is the number of cycles or steps.

We also calculated the circular standard deviation *σ*, which is a measure of the spread of the firing rate around the mean phase direction, and indicates where approximately 66% of the data lie (Drew and Doucet, 1991), as 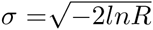. We calculated the Phase Selectivity Index (PSI). PSI is defined equivalently to the orientation selectivity index described by Mazurek et al. (2014),

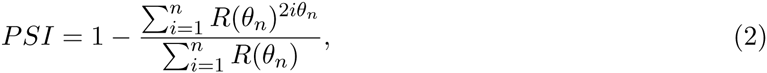

where *R*(*θ*) is the magnitude of the firing rate for any given angle *θ* = [0^*◦*^: 10^*◦*^: 360^*◦*^], for each stepping cycle.

### Mutual information

The Mutual Information between instantaneous firing rate and kinematic time-courses was computed using a continuous estimator based on the Kraskov, St¨ogbauer, and Grassberger (GSK) technique (Kraskov et al., 2004). We used the Matlab implementation of the GSK algorithm in the JIDT toolkit (Lizier, 2014). Firing rates and kinematics variables were computed as described above, and fed into the GSK algorithm, returning a mutual information value for each unit. For the step cycle calculation, the mutual information was computed between the firing rate and the phase angles of the Hilbert transform of the pitch velocity. Only periods during movement were considered, and mutual information was estimated for each cycle and then averaged. In this case, firing rate was computed every 5 ms and smoothed with a Gaussian filter of standard deviation 20 ms.

### Decoding

For every chosen experiment, the recorded cells’ spike trains were binned at 5 ms and then convolved with a Gaussian function (*σ* = 50, window width of 3*σ*) to obtain a time-course vector of instantaneous firing rates. The locomotion speed time course was convolved with the same Gaussian function. We considered all locomotion speeds *≤* 1 cm/s to be stationary; these were set to 0 cm/s. Both the firing rate and the locomotion speed time-courses were then normalised to obtain values between 0 and 1.

Only recording sessions with at least 8 units (resulting in inclusion of 6 sessions from 4 mice) were considered, in order to investigate the scaling of decoder performance with ensemble size. To decode, we used an optimal linear estimator (OLE) which weighted and linearly summed the instantaneous firing rate of each neuron in its ensemble, then rectified the summed output. We tested the incorporation of an additional offset term prior to rectification, but found that it did not improve performance on our dataset. The decoder was trained on 70% of each locomotion session, and tested on the remaining 30%. The OLE reconstruction is given by

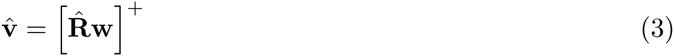

where 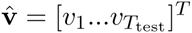 is the reconstructed locomotion speed time course, [.]^+^ denotes the rectification operator, 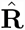 is a matrix whose columns consist of the firing rates **r**_*i*_ of each cell *i* for the *T*_test_ test time bins, and **w** is the linear estimator given by

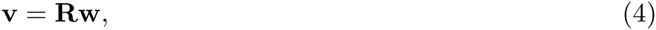

with **v** = [*v*_1_…*vT*_train_]^*T*^ being the measured speed time course vector for the *T*_train_ training data bins, and **R** a matrix whose columns are the firing rates for the training data, with the addition of a column of ones for the *y* intercept. Training the decoder by linear least squares regression is equivalent to solving this equation to find the optimal value of the estimator:

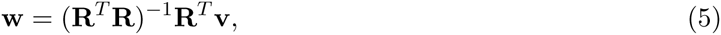

where **v** is a column vector containing the locomotion speed values for the training data. The estimated speed is half wave rectified to reflect the fact that only positive speed values are possible. We assessed decoding performance by computing the Pearson correlation coefficient between the actual and reconstructed locomotion speed time-courses, for the test data.

## Acknowledgements

This research was supported by the Biotechnology and Biological Sciences Research Council (BB/K001817/1), by a Royal Society Industry Fellowship (2011/R2) to SRS, by the M&B programme of Agenzia Regionale per il Lavoro, Sardegna, Italy (DR10-22462-1250/2011), and by the EU Marie Curie Initial Training Network NETT (EU FP7 289146). We thank Aldo Faisal for assistance in the development of the virtual reality setup.

## Supplementary Figures

**Figure 1 - figure supplement 1.**
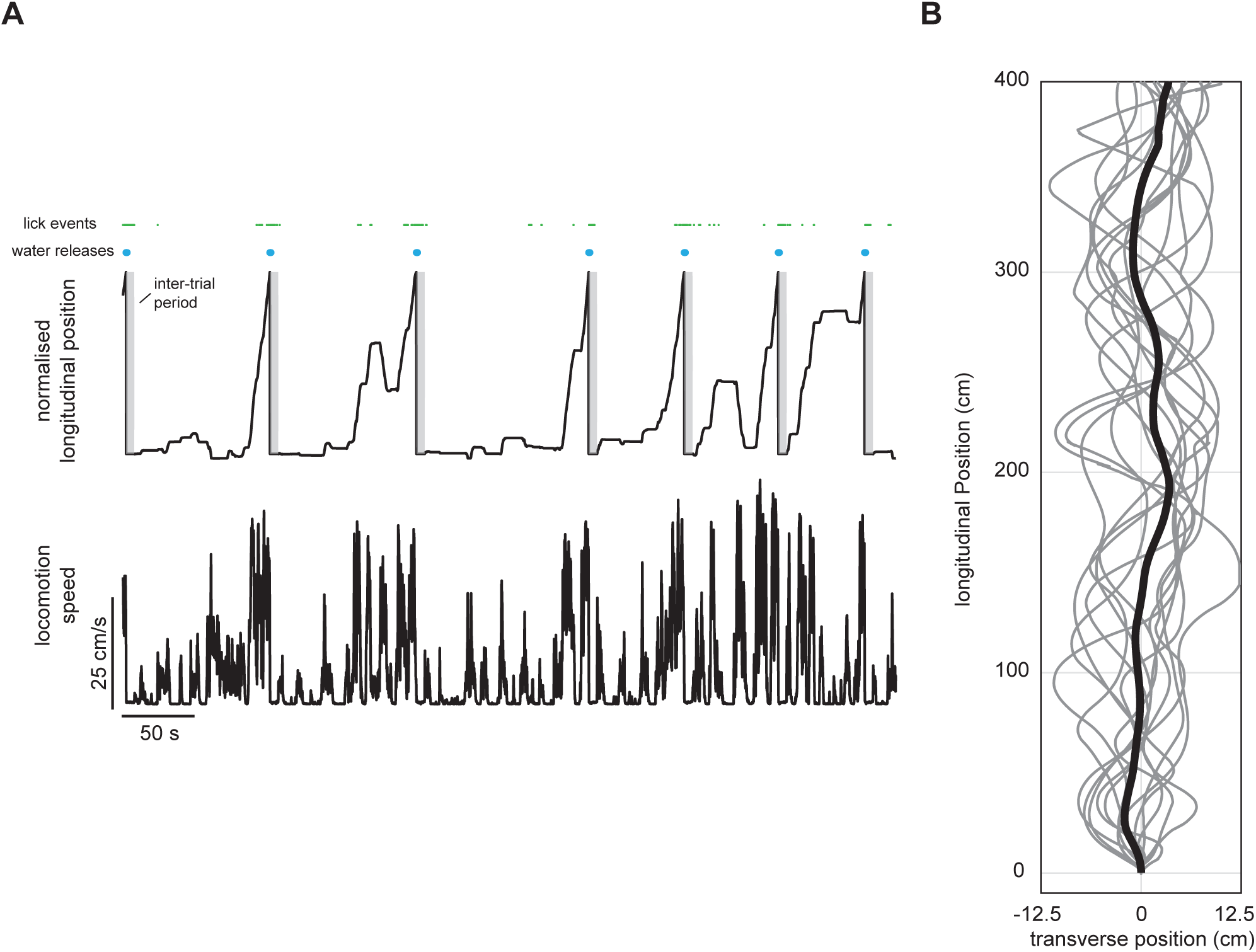
Behaviour in virtual reality environment. (**A**) Top: normalised longitudinal position of a mouse in the virtual corridor with respect to time during a few example trials with lick events and water pump activations. The mouse starts licking when it gets closer to the end as it expects the liquid reward for completing the task. Gray areas indicate inter-trial periods (duration = 6 seconds) during which there is no virtual reality projection and a black screen stimulus is presented. Bottom: speed signal as recorded by the computer mice positioned on the spherical treadmill. (**B**) A few example traces from the same experiment in A showing the trajectories in the virtual corridor. Please note the different scales along along the transverse and longitudinal directions. Black line indicate the mean trajectory of the trials shown.

**Figure 1 - figure supplement 2.**
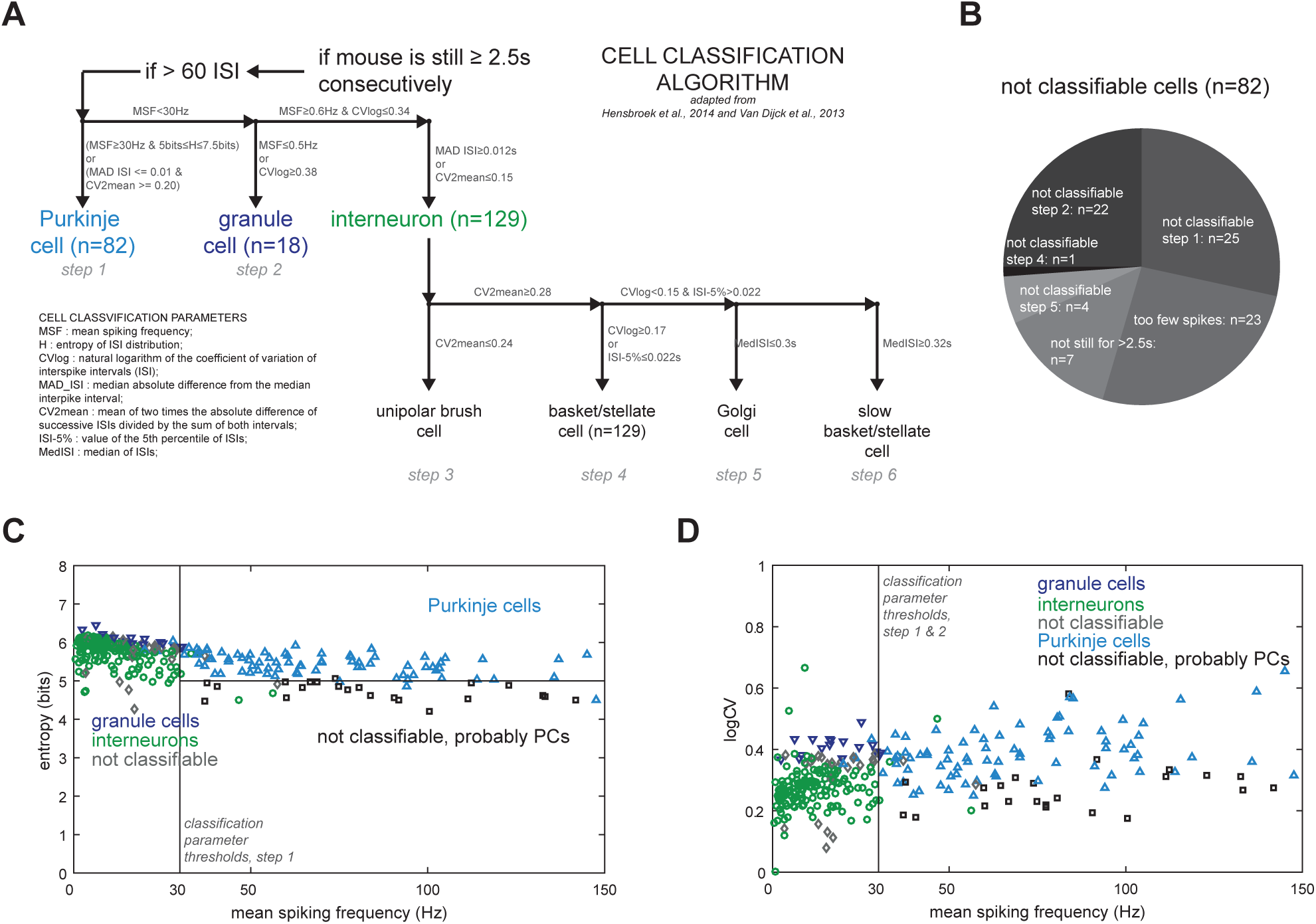
Identity classification of units. (**A**) Schematic of the classification algorithm based on the decision tree algorithm from Van Dijck et al., 2013 and Hensbroek et al., 2014. Purkinje, granule and interneuron cells are identified in the first three steps by using the parameters indicated on the figure. (**B**) Breakdown of the units that were not classifiable with reason and/or steps in which did not pass parameters thresholds. (**C**) Entropy and mean spontaneous spiking frequency of all units. Black lines show parameter thresholds for the first step of the classification algorithm. (**D**) Logarithm of the coefficient of variation of two consecutive ISIs and mean spontaneous spiking frequency of all units. Black line shows the parameter threshold for first classification step. Panel C and D are produced for comparison purposes with data shown by Van Dijck et al., 2013 in their research article.

**Figure 2 - figure supplement 1.**
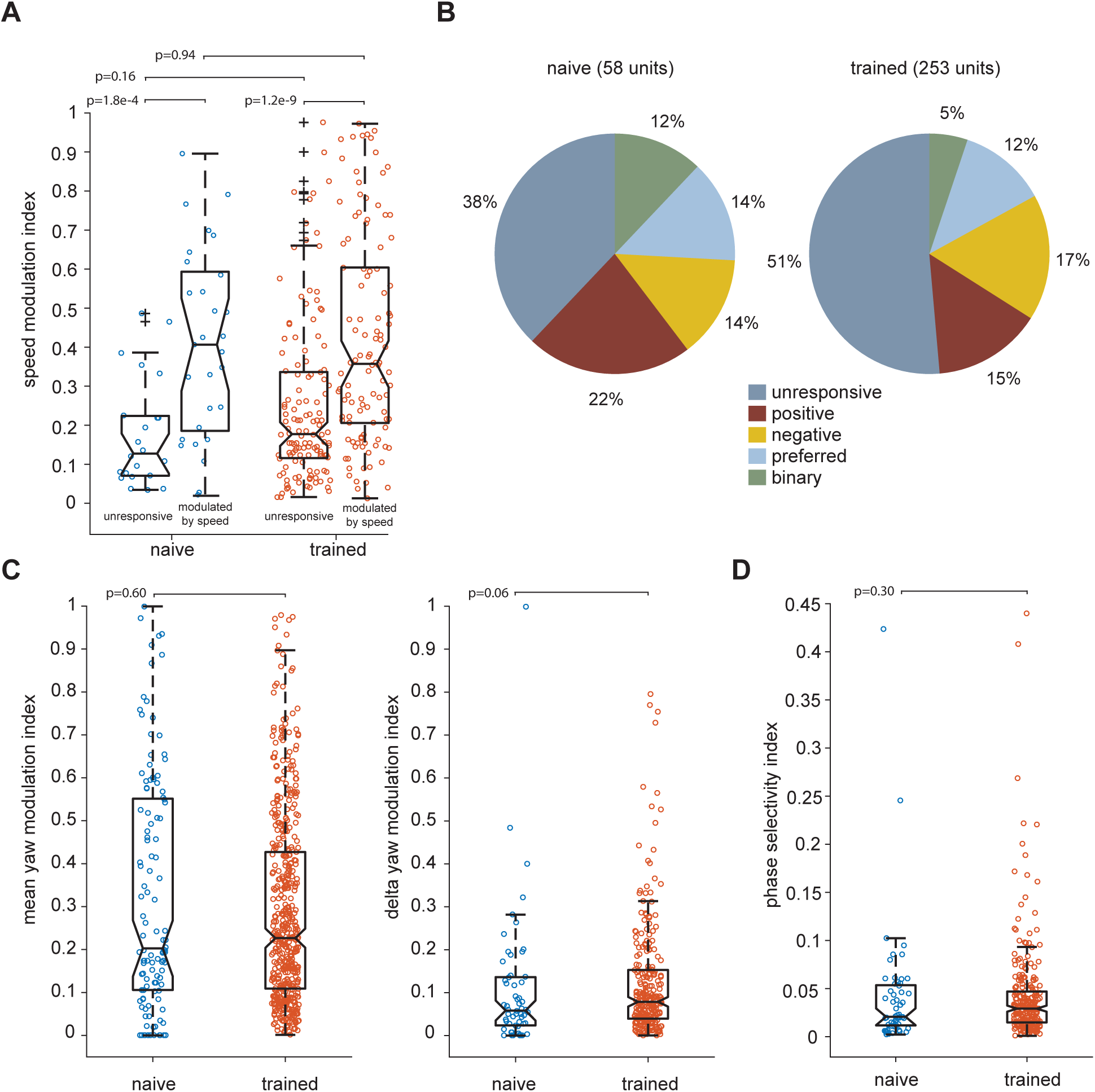
No differences in the neural populations recorded in na¨ıve or trained animals. (**A**) The speed modulation indexes are not significantly different between na¨ıve (total number of units=58, units modulated by speed=29) or trained animals (total number of units=253, units modulated by speed=110) for unresponsive or speed modulated units. (**B**) Classification of speed-response profiles in the na¨ıve versus trained conditions: the percentages of each type is similar in both conditions. (**C**) Both the mean and the absolute difference between the modulation indexes between CW and CCW yaw tuning curve (delta yaw modulation index) are not significantly different. (**D**) No significant difference is found in the phase selectivity index (PSI) in the na¨ıve versus the trained animals.

**Figure 2 - figure supplement 2.**
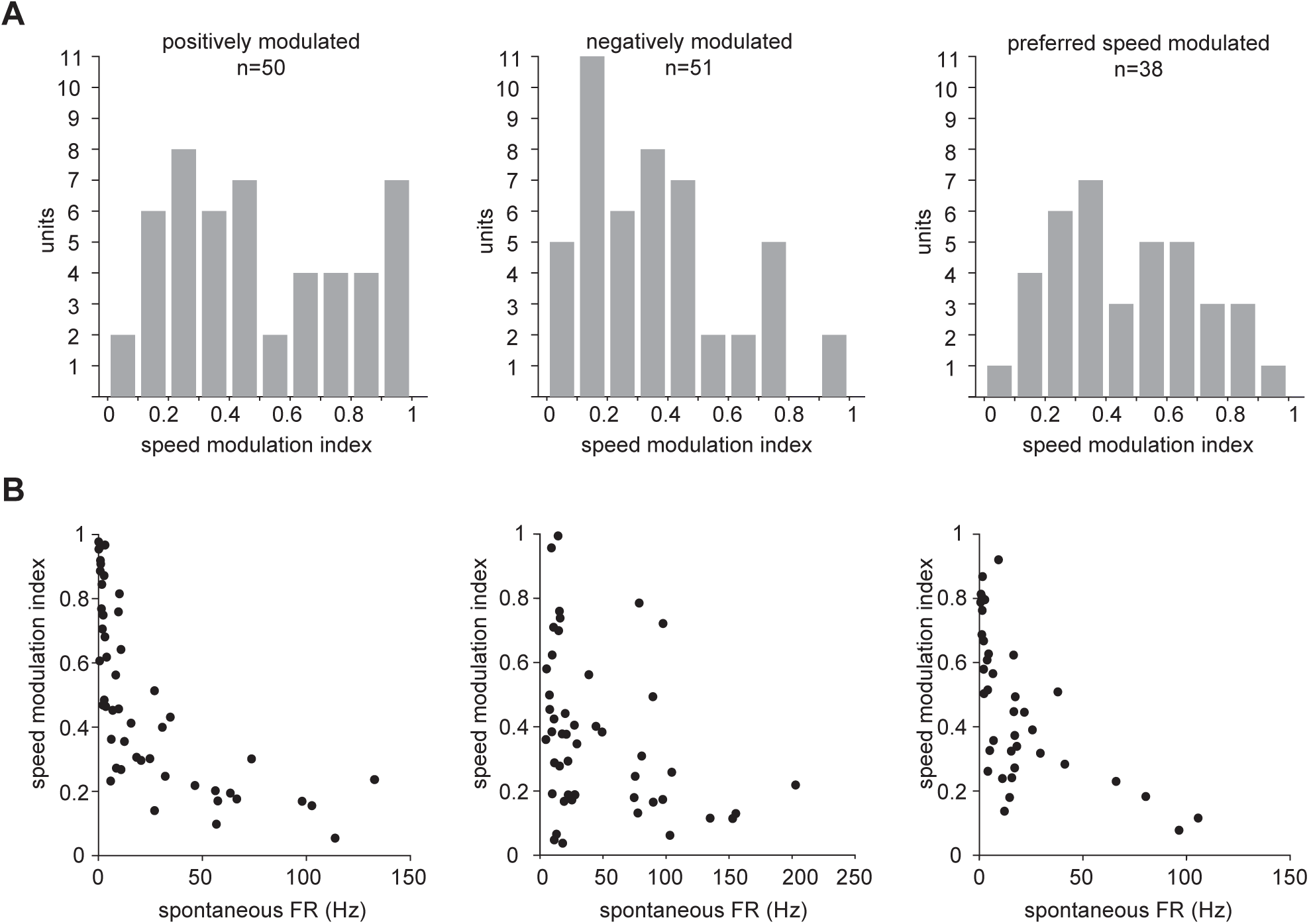
Modulation of speed tuned units. (**A**) Modulation indexes of all units significantly modulated by speed, divided by response type. (**B**) Modulation indexes shown as a function of spontaneous firing rate (i.e. during stationary periods), divided by response type.

**Figure 2 - figure supplement 3.**
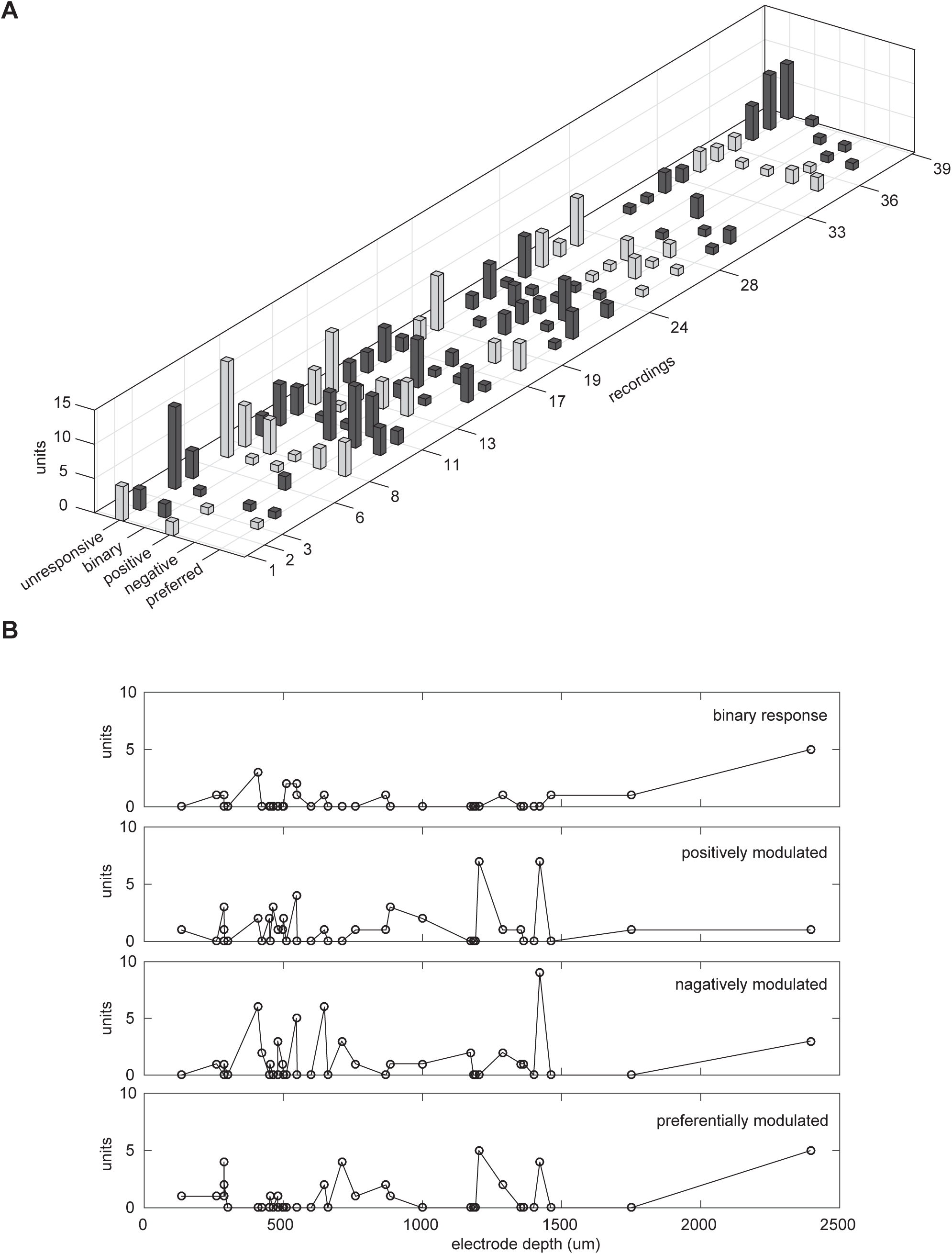
Count of units per recording. (**A**) Count of units for every recording used in this study classified according to their response profile to locomotion speed. Black and white bars are used only to help distinguishing between consecutive animals; ticks on the recordings axes sign the switch to different animal. (**B**) Poll of units responsive to movement with respect to the depth of each recording.

**Figure 3 - figure supplement 1.**
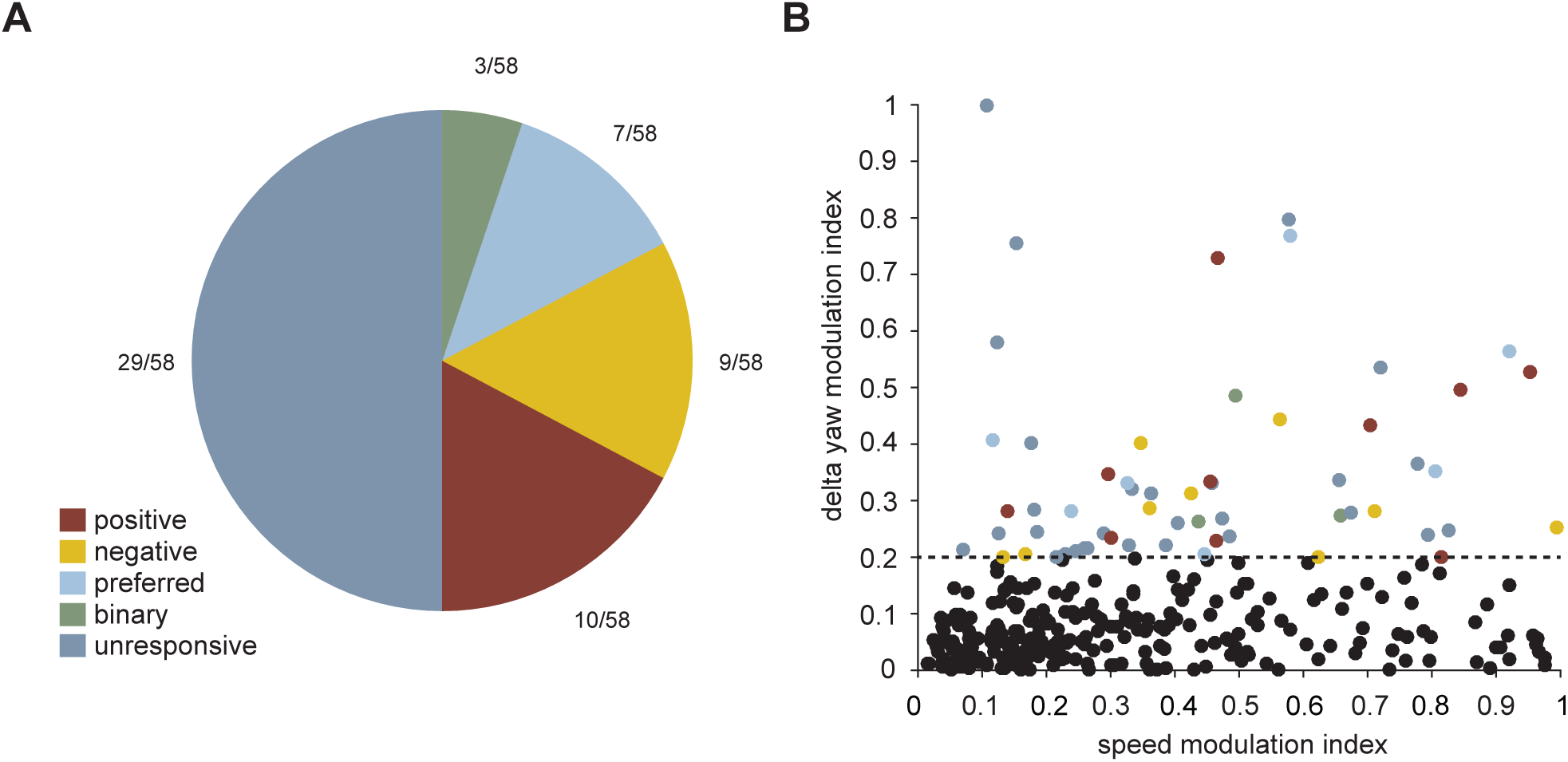
Yaw tuning responses with respect to speed tuning responses.(**A**) Out of the 58 units with an absolute difference in modulation indexes between CW and CCW yaw tuning curve (delta yaw modulation index) larger than 0.2, more than half are not modulated by speed, while the rest are evenly spread amongst the other response categories. (**B**) A large delta yaw modulation index is not related to the speed modulation index (n=311). The dashed line is the threshold that separates the 58 units represented in A.

**Figure 6 - figure supplement 1.**
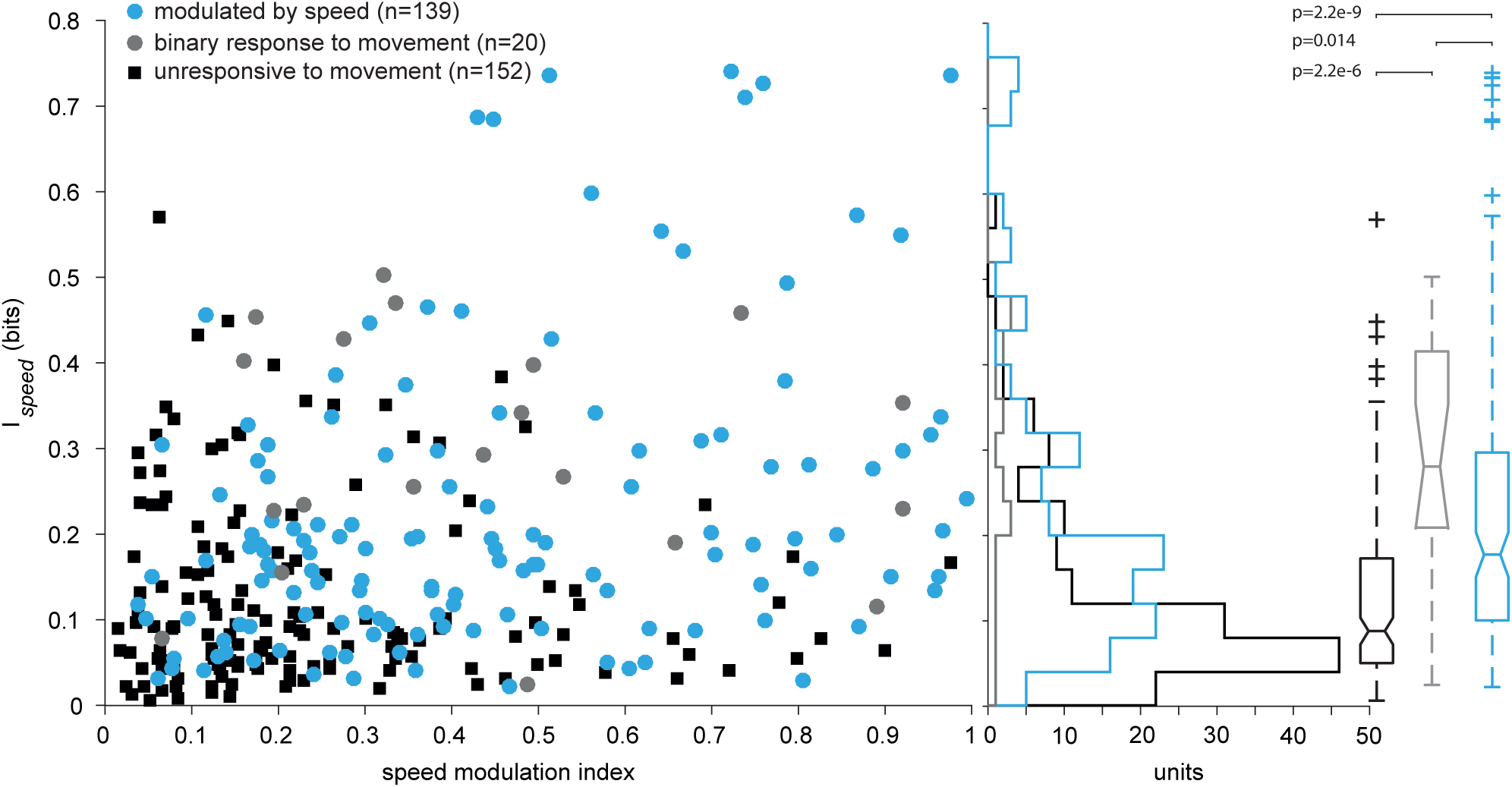
Mutual information versus speed modulation index. Mutual information of locomotion speed computed at zero lag as a function of the modulation index for speed modulated for all units. P values of Mann-Whitney U-test between the three distributions.

